# Bacteriostatic antibiotics drive bacterial death by reshaping competitive interactions

**DOI:** 10.64898/2026.07.22.739985

**Authors:** Valentin Rodriguez, Justine Gillard, Katharina Müller, Katia Villion, Yanis Alcivar, Chloé Virolle, Camille V. Goemans

## Abstract

Protein synthesis inhibitors, including macrolides and tetracyclines, are classically defined as bacteriostatic because they arrest bacterial growth without directly causing death. Yet their effects vary widely across bacterial species and communities, suggesting that additional mechanisms shape their ecological impact. Here we show that the human gut commensal *Escherichia coli* ED1a kills the laboratory strain BW25113 specifically when protein synthesis is inhibited. We find that this killing is mediated by colicin K, a bacteriocin encoded on a plasmid carried by ED1a alongside its cognate immunity factor. Because ED1a itself is far more sensitive to protein synthesis inhibitors than BW25113 (surviving antibiotic exposure roughly 1,000-fold less efficiently), we expected antibiotic treatment to favour BW25113 in co-culture. Instead, we found that antibiotic exposure suppresses BW25113 below the population density required to coexist with colicin-producing ED1a, triggering colicin-dependent elimination of the sensitive strain, ultimately leading to the collapse of both populations. Screening a library of 1,085 *E. coli* isolates revealed that this antibiotic-induced, toxin-mediated killing is not unique to the ED1a–BW25113 pair but is a widely conserved interaction across natural strain combinations. Thus, a nominally bacteriostatic antibiotic can indirectly drive bacterial death by reshaping competitive dynamics between strains rather than by any direct bactericidal action. These findings reveal a previously unrecognized route by which protein synthesis inhibitors influence bacterial competition, and offer a framework for understanding how clinically important bacteriostatic antibiotics can eliminate bacterial populations and restructure microbial communities.

## Introduction

Over the past century, antibiotics have become a cornerstone of modern medicine, dramatically increasing life expectancy and transforming the treatment of infectious diseases. However, their widespread and often indiscriminate use, combined with the remarkable adaptive capacity of bacteria, has driven the emergence of strains resistant to virtually all clinically available antibiotics. As a result, the World Health Organization has recognized antibiotic resistance as a major global health threat, underscoring the urgent need for new antimicrobial strategies and alternative approaches to combat resistant pathogens. In parallel, a comprehensive understanding of how existing antibiotics affect diverse bacterial species and communities remains essential to optimize therapeutic outcomes and mitigate the development of resistance.

Antibiotics are typically classified as either bactericidal (killing bacteria) or bacteriostatic (preventing bacterial growth without killing). While this distinction remains prevalent in textbooks and clinically important^1^, we^2^ and others^1,3–8^ have increasingly challenged this dogma. We previously studied macrolides and tetracyclines, protein synthesis inhibitors associated with strong dysbiosis of the human gut microbiota^9–13^. These antibiotics have been used clinically for decades, and their targets and modes of action have been extensively characterized, primarily in model organisms and common pathogens. It is textbook knowledge that, with the exception of aminoglycosides, protein synthesis inhibitors block translation reversibly and are therefore classified as bacteriostatic. However, we previously showed that macrolides and tetracyclines are in fact bactericidal against several human gut bacterial strains^2^, and that in a synthetic community, antibiotics that are bacteriostatic on some strains but bactericidal on others induce substantial shifts in community composition, potentially explaining the long-term dysbiosis observed in humans and mice after treatment^9–13^.

More recently, we observed that the *E. coli* gut commensal ED1a shares the same minimal inhibitory concentration (MIC) for protein synthesis inhibitors as the laboratory strain BW25113, yet survives antibiotic treatment over 1,000-fold less efficiently^2,14^. In other words, despite identical antibiotic sensitivity as measured by MIC, ED1a is effectively eliminated by treatment while BW25113 survives. Even more strikingly, when the two strains were co-cultured in the presence of a protein synthesis inhibitor, both populations were eliminated, suggesting that ED1a exerts a toxic effect on BW25113 that is unmasked specifically under antibiotic pressure.

Here, we set out to understand how bacteriostatic protein synthesis inhibitors trigger the killing of BW25113 in the presence of ED1a. We show that this competitive outcome is driven by the bacteriocin colicin K. ED1a harbours a small plasmid encoding both colicin K and its cognate immunity protein, allowing it to selectively eliminate susceptible competitors such as BW25113. Critically, this antagonism is conditional on antibiotic treatment, which suppresses target populations to densities low enough to permit colicin-mediated clearance. Moreover, we show that this mechanism drives clearance of BW25113 from a synthetic community of 12 gut bacterial members that also includes ED1a, when exposed to protein synthesis inhibitors. Screening a library of 1,085 *E. coli* isolates, we find that this antibiotic-induced, toxin-mediated killing is not restricted to the ED1a–BW25113 pair but is a widely conserved interaction across natural strain combinations. Together, our findings reveal that antibiotics can indirectly shape bacterial competition by modulating toxin-mediated interactions between strains. This work provides a mechanistic framework linking antibiotic exposure to shifts in microbial community composition, and shows how bacteriocins encoded within bacterial genomes can synergize with antibiotics to shape ecological outcomes. More broadly, our study uncovers a previously unappreciated mechanism of antibiotic action, advancing our understanding of how commonly used bacteriostatic antibiotics eliminate bacterial populations and reshape microbial communities.

## Materials and methods

### Strains, plasmids and growth conditions

All strains (Table S1) and plasmids (Tables S2) used in this study appear in the supplemental information. All strains were grown in LB (Lennox, Sigma Aldrich) at 37 °C unless indicated otherwise. All experiments were performed at mig-log phase (OD_600nm_∼0.3-0.4) after diluting an overnight culture and growing the subculture to the aforementioned OD. All BW25113 knock-out strains derive from the Keio collection^15^. All clonings were perfomed by Gibson Assembly. The primers were designed with the Gibson Assembly tool from SnapGene (version 8.0.3). The assemblies were performed using the NEBuilder^®^ HiFi DNA Assembly from NEB and following the manufacturer’s protocol.

### Minimal Inhibitory Concentration (MIC) measurement

Overnight cultures were diluted to OD_600mn_ = 0.01 and cultured with antibiotics at eight concentrations in a twofold dilution gradient, in three technical replicates in microtiter plates (U-bottomed 96-well plates; Greiner Bio-One, 268200). Plate were sealed with breathable membrane (Breathe-Easy; Sigma-Aldrich, Z380059-1PAK) and grown at 37 °C with continuous shaking (850 rpm; orbital microplate shaking). The OD_600nm_ was measured every 30 min for 24 h using the Agilent BioTek Synergy H1 plate reader. All MIC tests were performed in a total volume of 100 µl per well.

### Survival assays

Overnight cultures were diluted 1:100 in fresh LB in the morning. Once the OD_600nm_ reached ∼0.3-0.4 (or a different OD if specified), the cultures were treated with 5 x MIC of the given antibiotic (final concentration is 20 µg/ml of chloramphenicol). A 100 µl of each culture was collected and serial-diluted in PBS using a 10-fold dilution step in microtiter plates. 3 µl of each dilution were then spotted on LB-Agar and incubated overnight at 37°C. After 5h of treatment (unless specified otherwise), the same dilution and spotting steps were repeated. The next day, colony forming units (CFU) were counted and the survival percentages calculated by dividing the number of surviving cells at time 5 h by the number of surviving cells at time 0, multiplied by 100.

### Lysis assays – individual cultures

Overnight cultures were diluted 1:100 in fresh LB in the morning. Once the OD_600nm_ reached ∼0.3-0.4, the cultures were treated with 5 x MIC of the given antibiotic (20 µg/ml of chloramphenicol) and transferred in a microtiter plate to measure lysis with a plate reader, or in glass tubes/flasks when a higher volume was necessary. The microtiter plates were sealed (Breathe-Easy; Sigma-Aldrich, Z380059-1PAK) and OD_600nm_ was measured in an Agilent LogPhase600 plate reader for max 24 h, with a 20 min interval. When measured in tubes or flasks, 1 ml of culture was collected in a plastic 1 cm-cuvette at each time point and OD_600nm_ measured using a Biochrom Libra S60 UV/Vis spectrophotometer.

### Supernatant preparation and tests

ED1a (or any other specified strain) was grown overnight. In the morning, the culture was diluted 1:100 and grown until reaching an OD_600nm_ of 0.3, at which point the antibiotic was added (at 5 x MIC) or not, depending on the experiment. When the supernatant was prepared without antibiotic, the culture was further grown until reaching an OD_600nm_ of ∼3. When the supernatant was prepared with antibiotic, the culture was further incubated for 5h, without visible increase in OD. The cultures were then centrifuged (4000 rpm, 10 min, RT), the supernatant collected and filtered on a PVDF filter, 0.22 µM. The supernatants were kept at 4°C for max 1 week. When specified, DnaseI (2 U/µl; 1 µl per ml), RnaseA (10 mg/ml; 1 µl per ml) or Proteinase K (20 mg/ml; 3 µl per ml) were added to the supernatant and incubated at 37°C for an hour.

### Supernatant fractionation and mass spectrometry

ED1a was grown overnight. In the morning, the culture was diluted 1:100 in 1L fresh LB and grown until reaching an OD_600nm_ of 3. The culture was then centrifuged at 4000 rpm for 20 min, the supernatant collected and filtered on a PVDF filter, 0.22 µM. The supernatant was then concentrated 400X using a 30kDa cut-off Amicon Ultra filter (Millipore) and loaded on a Superdex 200, 10/300 (using the AKTAPure system), previously equilibrated with 1x PBS. The resulting fractions were collected and tested for killing activity (see survival tests). Fractions with and without killing activity tested by mass spectrometry (Proteomics Core Facility at EPFL).

A volume of 300 uL per fraction was mixed with 2x SDS buffer (4 % SDS, 200 mM Tris-HCl, pH 8.0 supplemented with protease inhibitor) in a 1:1 volume ratio, boiled at 90°C for 10 minutes and concentrated to a final volume of 75 µL using vacuum centrifugation. The concentrated fractions were digested following the S-Trap™ micro spin column digestion protocol (Protifi) with minor adaptation. Briefly, samples were reduced for 10’ at 95°C in 10 mM TCEP and alkylated by incubation for 30 minutes in 20 mM Iodoacetamide (Sigma-Aldrich) at room temperature. After acidification using 27.5 % phosphoric acid, the samples were diluted in binding buffer in a 1:6 volume ratio and loaded on the columns. Sample cleaning and digestion were performed according to the manufacturer recommendation using a 50 mM Ammonium bicarbonate-based digestion buffer overnight. After peptide recovery by a first centrifugation step, digested samples were further eluted sequentially using 40 μl 50 mM AB, 0.2 % formic acid and 50 % acetonitrile. Finally, peptides were desalted on SDB-RPS StageTips^16^ and dried by vacuum centrifugation until further use.

Samples were resuspended in 2% acetonitrile (Biosolve), 0.1% FA and nano-flow separation was performed using a Vanquish Neo nano UPLC system (Thermo Fischer Scientific) on-line connected with an Orbitrap Fusion Lumos Tribrid Mass Spectrometer (Thermo Fischer Scientific). A capillary precolumn (Acclaim Pepmap C18, 3 μm-100Å, 2 cm x 75μm ID) was used for sample trapping and cleaning. A 50 cm long capillary column (75 μm ID; in-house packed using ReproSil-Pur C18-AQ 1.9 μm silica beads; Dr. Maisch) was then used for analytical separations at 250 nl/min over 150 min biphasic gradients. Acquisitions were performed through Top Speed Data-Dependent acquisition mode using a cycle time of 1 second. First MS scans were acquired with a resolution of 240K (at 200 m/z) on the orbitrap and the most intense parent ions were selected and fragmented by High energy Collision Dissociation (HCD) with a Normalized Collision Energy (NCE) of 30% using an isolation window of 0.7 m/z. Fragmented ions were acquired on the ion trap with a maximum injection of 20 ms and selected ions were then excluded for the following 20 s.

Raw data were processed using Sequest HT and MS Fragger^17^ in Proteome Discoverer v.2.5 against the ED1a Uniprot database supplemented with the common MaxQuant contaminant entries. A minimum of two peptides were required for protein identifications. Enzyme specificity was set to trypsin and a minimum of six amino acids was required for peptide identification. Tolerance values of 20 ppm at parent level and 0.3 Da at fragment ones were used. Up to two missed cleavages were allowed and a 1% false discovery rate (FDR) cut-off was applied both at peptide and protein identification levels. For the database search, carbamidomethylation (C) was set as fixed modifications whereas oxidation (M) was considered as a variable one.

### Small plasmid isolation

To isolate small plasmids from ED1a, we took advantage of an ED1a Tn-library (with Kan^R^) built in the lab, with the hope that some transposons (and their Kan-resistance cassette) landed in the plasmids. The rationale was that small plasmids typically do not carry antibiotic resistance genes for their selection, so they are typically quite difficult to isolate. The ED1a Tn-library plasmids were isolated using a miniprep kit (QIAprep Spin Miniprep kit, QIAGEN). After quantifying the product by nanodrop, the mixture of isolated plasmids was transformed into BW25113 competent cells and selected on kanamycin (50 µg/ml), reaching 14 colonies, each carrying a single plasmid. These colonies were further grown in LB+ kanamycin (50 µg/ml), and their respective plasmid isolated using a miniprep kit (QIAprep Spin Miniprep kit, QIAGEN). The plasmids were sequenced (Microsynth – Oxford Nanopore Sequencing – FullPlasmidSeq), and their sequences compared to existing plasmids (NCBI Blasts).

### Immunity proteins overexpression

Overnight cultures of BW25113 strain containing plasmids overexpressing colK-IP (pUC18-colK-IP) and pest-IP (pUC18-pestIP), and the corresponding empty vector, were diluted 1:100 with addition of 1 mM IPTG for induction. After 1.5h, cells were centrifuged and resuspended in ED1a supernatant and chloramphenicol (final concentration of 20 µg/ml, 5 x MIC) and incubated for 5h. CFU spotting and counting was performed as in *Survival assays* above.

### Small plasmid elimination from ED1a

For this part, we used an *E. coli* MFDpir donor strain carrying a miniTn7 transposon with an arabinose-inducible copy of *ddmDE* on a plasmid e.g. pGP704-TnAraC-ddmDE^18^, *E. coli* MFDpir donor with helper plasmid pUX-BF13 and our recipient strain ED1a.

All 3 strains were grown overnight in LB at 37°C. Donor strain cultures were supplemented with 100 µg/mL Amp (plasmid maintenance) and 0.3 mM DAP (MFDpir strain is DAP-dependent). 0.6 mL each O/N culture were harvested (20,000 x g; RT; 3min), washed once with 0.6 mL PBS, and resuspended in final volume of 0.3 mL PBS. Cultures were mixed 1:1:1 and vortexed 5-10s. 150 µL of mix was spotted onto a LB+ 0.3mM DAP plate, and incubated without inverting at 37°C for ∼6h. The material from the spot was picked and spread out on a LB+Gent25 plate, and incubated at 37°C, O/N. Independent colonies were patched onto LB+Gent25 **and** LB+Amp100 plates to identify and remove unwanted AmpR clones in which the complete plasmid has integrated onto the recipient chromosome. Several clones were checked by colony PCR to verify insertion, using primers NotI-90-outwards (GCTTTAGCCATAACAAAAGTCCAG) that binds in the transposon and Ec_glmS-end-out (GCCGCATGTGGAAGAGGTGATTGC) that binds the genome of *E. coli*. Culture were then grown overnight with or without arabinose (0 or 0.4%) and miniprepped (QIAGEN QIAprep Spin Miniprep Kit)to measure the amount of remaining plasmids.

### BW25113 treatment with ED1a supernatant and mass spectrometry

Five overnight cultures of BW25113 were each diluted 1:100 in 10 ml fresh LB and grown until reaching OD_600nm_ = 0.3. Next, each 10 ml culture was split into two and centrifuged at 4000 rpm, 10 min, RT to collect the pellets. For each replicate, one pellet was directly resuspended with PBS while the other one was resuspended in ED1a supernatant (no antibiotic) for 30 min. Pellets were then all washed twice with PBS and resuspended in 50 µl of lysis buffer (100 mM Tris pH7.5, 2% SDS, protease inhibitors). The samples were incubated for 10 min at 90°C with shaking (500 rpm), and further sonicated 5x 30 sec. Next, all samples were centrifuged at top speed to remove cell debris, and the supernatant was collected and frozen at -20°C.

Mass spectrometry-based proteomics-related experiments were performed by the Proteomics Core Facility at EPFL. Proteins were digested by filter aided sample preparation (FASP^19^) with minor modifications. Proteins (20 μg) were deposited on top of washed and conditioned Microcon®-30K devices (Merck AG, Zug, Switzerland). Samples were centrifuged at 9400×g, at 20°C for 30 min or until complete dryness. All subsequent centrifugation steps were performed using the same conditions. Two washing steps were performed using 200 μL urea solution (8 M Urea, 100 mM Tris-HCl pH 8). Reduction was performed by adding 100 μL of 10 mM Tris(2-carboxy)phosphine (TCEP) in urea solution on top of filters followed by 60 min incubation time at 37°C with gentle shaking and light protection. Reduction solution was removed by centrifugation and two washing steps with 200 μL urea solution. Then, alkylation was performed by adding 100 μL of 40 mM chloroacetamide (CAA) in urea solution and incubating the filters at 37°C for 45 min with gentle shaking and protection from light. The alkylation solution was removed by centrifugation followed by two washing steps with 200 μL of urea solution. Finally, two additional washing steps using 200 μL of 5 mM Tris-HCl pH 8 were performed to condition the filters for digestion. Proteolytic digestion was performed overnight at 37°C by adding 100 μL of a combined solution of Endoproteinase Lys-C and Trypsin Gold in an enzyme/protein ratio of 1:50 (w/w) prepared in 5 mM Tris-HCl and 10 mM CaCl2 on top of filters. The resulting peptides were recovered by centrifugation and two subsequent elution with 50 μL of 4% trifluoroacetic acid. Finally, the recovered peptides were desalted on SDB-RPS StageTips^20^ and dried by vacuum centrifugation prior to LC-MS/MS injections.

Samples were resuspended in 2% Acetonitrile, 0.1% Formic acid and nano-flow separations were performed on a Vanquish Neo nano UPLC system (Thermo Fischer Scientific) on-line connected with an Exploris 480 Orbitrap Mass Spectrometer. A capillary precolumn (Acclaim Pepmap C18 ; 3μm-100Å ; 2cm x 75μm ID) was used for sample trapping and cleaning. Analytical separations were performed at 250nl/min over a 150min. biphasic gradients on a 50cm long in-house packed capillary column (75μm ID; ReproSil-Pur C18-AQ 1.9μm silica beads; Dr. Maisch). Mass spectrometry data were acquired on an Orbitrap Exploris mass spectrometer (Thermo Fisher Scientific) operated in data-independent acquisition (DIA) mode. Full MS1 scans were acquired in the Orbitrap over an m/z range of 420-680 at a resolution of 60000 (at m/z 200), with a normalized automatic gain control (AGC) target of 300% and a maximum injection time set at auto. MS1 scans were followed by sequential DIA MS2 scans covering an m/z range of 430-670, using non-overlapping 6.6 m/z isolation windows resulting in 36 DIA windows per cycle. Fragmentation was performed by higher-energy collisional dissociation (HCD) using a normalized stepped collision energy of 22,26 and 30. MS2 spectra were acquired in the Orbitrap at a resolution of 45000 (at m/z 200) over an m/z range of 200-1800, with a normalized AGC target % at 3000.

Raw data files were analysed with DIA-NN^21^ using library free search against the ED1a and BW25113 uniprot database^22^. Results were filtered at 1% FDR. Search parameters included: minimum fragment m/z at 200, maximum fragment m/z at 1800, N-terminal methionine excision enabled, *in silico* digest with cuts at K* and R*, maximum number of missed cleavages set to 1, minimum peptide length set to 7, maximum peptide length set to 30, minimum precursor m/z at 300, maximum precursor m/z at 1800, minimum precursor charge set to 1, maximum precursor charge set to 4. Carbamidomethylation (C) was set as a fixed modification, whereas oxidation (M) as a variable modification. Maximum number of variable modifications was set to 2 and Match between runs (MBR) was enabled. The Unrelated runs option was checked. Resulting DIANN report was converted to MSstats format using MSstatsConvert (v1.18.1) and analyzed in R with MSstats^23^. Differential protein expression analysis was performed using the linear mixed-effects model followed by the Benjamini– Hochberg procedure. Adjusted P value < 0.05 (and |log2 fold change| > 1) were considered as significant.

### Microscopy (microfluidics)

An overnight culture of BW25113 was diluted 1:100 and grown for 2h. In the meantime, the microfluidics plate (ONIX, CellASIC®) was prepared by removing the PBS and adding the LB or ED1a supernatant, containing or not Cm (20 µg/ml), and incubated on the microscope to heat to 37°C. After 2h, cells were treated or not with Cm (20 µg/ml) and transferred to the microfluidics plate for imaging. 100 µl of the culture was loaded into the B04A microfluidic chamber. The nutrient supply was maintained at 3 psi, and the temperature was maintained throughout the imaging process using temperature-controlled stage-top chamber (H301-K-Frame; OKOLAB) set to 37 °C with an accompanying objective heater. Cells were imaged automatically every 20 min for 7 hours. Imaging was conducted using Zeiss Axio Observer Z1 epifluorescence microscope, equipped with AxioCam MRm camera and controlled by Zeiss Zen Software (v.2.6 blue edition). Images were captured using a Plan-Apochromat 100x/1.4-NA Ph3 oil objective, and illumination was provided by an HXP120 lamp. Image analysis was performed using Fiji.

### Lysis assays – screen

Deep-well plates were filled with 1mL of LB in each well and pre-warmed to 37 °C for 30 min before transfer of 10 µL of overnight culture of each *E. coli* isolate. Plates were grown at 37 °C for 6h shaking at 180 rpm, and centrifuged to collect supernatants. If applicable, supernatants were treated with Proteinase K (20 mg/ml; 3 µl per ml) for 1 hour at 37 °C.

An overnight culture of BW25113 cells was diluted 1:100 in the morning and grown in a flask at 37 °C until reaching an OD_600nm_∼0.6. Cells were then centrifuged and resuspended in 10X less volume with 50 X MIC of chloramphenicol (200 µg/ml). 15 µL of this culture was then dispensed in each well and mixed with 135 µL of each *E. coli* isolate supernatant. The microtiter plates were sealed (Breathe-Easy; Sigma-Aldrich, Z380059-1PAK) and OD_600nm_ was measured in an Agilent LogPhase600 plate reader for max 24 h, with a 20 min interval.

### Community experiment

All subsequent experiments were performed inside an anaerobic chamber. Strains (*B. fragilis, B. uniformis, B. thetaiotaomicron, B. ovatus, B. caccae, P. vulgatus, P. copri, F. nucleatum, C. bolteae, E. rectale, R, intestinalis, E. coli* ED1a and *E. coli* BW25113-Kan^R^) were grown in 5 ml of MGAM medium and incubated at 37 °C overnight. After the first overnight, all strains were diluted again 1:100 in 5 mL fresh MGAM and grown a second overnight. Strains were then mixed 1:1:1:… in terms of OD to assemble the communities, and diluted 100x in fresh MGAM. Community 1 contains all the strains except for ED1a, while community 2 contains all the strains including ED1a. Communities were grown for 1h before chloramphenicol (20 µg/ml) was added for 5h of treatment. Communities were spotted for CFU (as described in *Survival assays* above) at t0 and t5 on (i) MGAM plates, for all community detection, (ii) LB plates incubated outside the anaerobic chamber, for all *E. colis* detection, (iii) LB+Kan plates incubated outside the anaerobic chamber, for *E. coli BW25113* detection.

## Results

### A toxic protein in the supernatant of ED1a kills BW25113 upon chloramphenicol treatment

We previously observed that ED1a, an *E. coli* human gut commensal was killed by *bacteriostatic* protein synthesis inhibitors, while the lab strain BW25113 survived the bacteriostatic treatment^2,14^ (**Fig. 1A**). Surprisingly, we observed that mixing the two strains during treatment (here, chloramphenicol) led to the elimination of both strains, suggesting that the presence of ED1a in the culture turns chloramphenicol into a bactericidal antibiotic for BW25113 (**Fig. 1A**). To determine whether this effect was direct or indirect, we isolated ED1a supernatant and tested its effect on BW25113. While the supernatant or chloramphenicol alone did not show decreased survival of BW25113, their combination showed high toxicity (**Fig. 1B**), accompanied by strong lysis (**Fig. 1C**). To determine the nature of the killing, the supernatant was pre-treated with proteinase K, DnaseA or RnaseI and combined with chloramphenicol before exposure to BW25113. While the supernatants treated with DnaseA or RnaseI retained their killing activity, the supernatant treated with proteinase K completely lost its toxicity (**Fig. 1B**), suggesting the presence of a toxic protein in the supernatant of ED1a. In parallel, we explored whether specific BW25113 surface proteins were necessary for the toxic activity of the ED1a supernatant. To address this, we selected a panel of 26 proteins, all of which are present on BW25113 surface or known to function as importers. We used the 26 corresponding deletion mutants and tested whether they lysed when exposed to ED1a supernatant and chloramphenicol. From the 26 mutants tested, only three survived the treatment: Δ*tsx*, Δ*tonB* and Δ*tolC* (**Fig. 1D**), suggesting that those could be involved in the import of the toxic protein.

**Figure 1.**
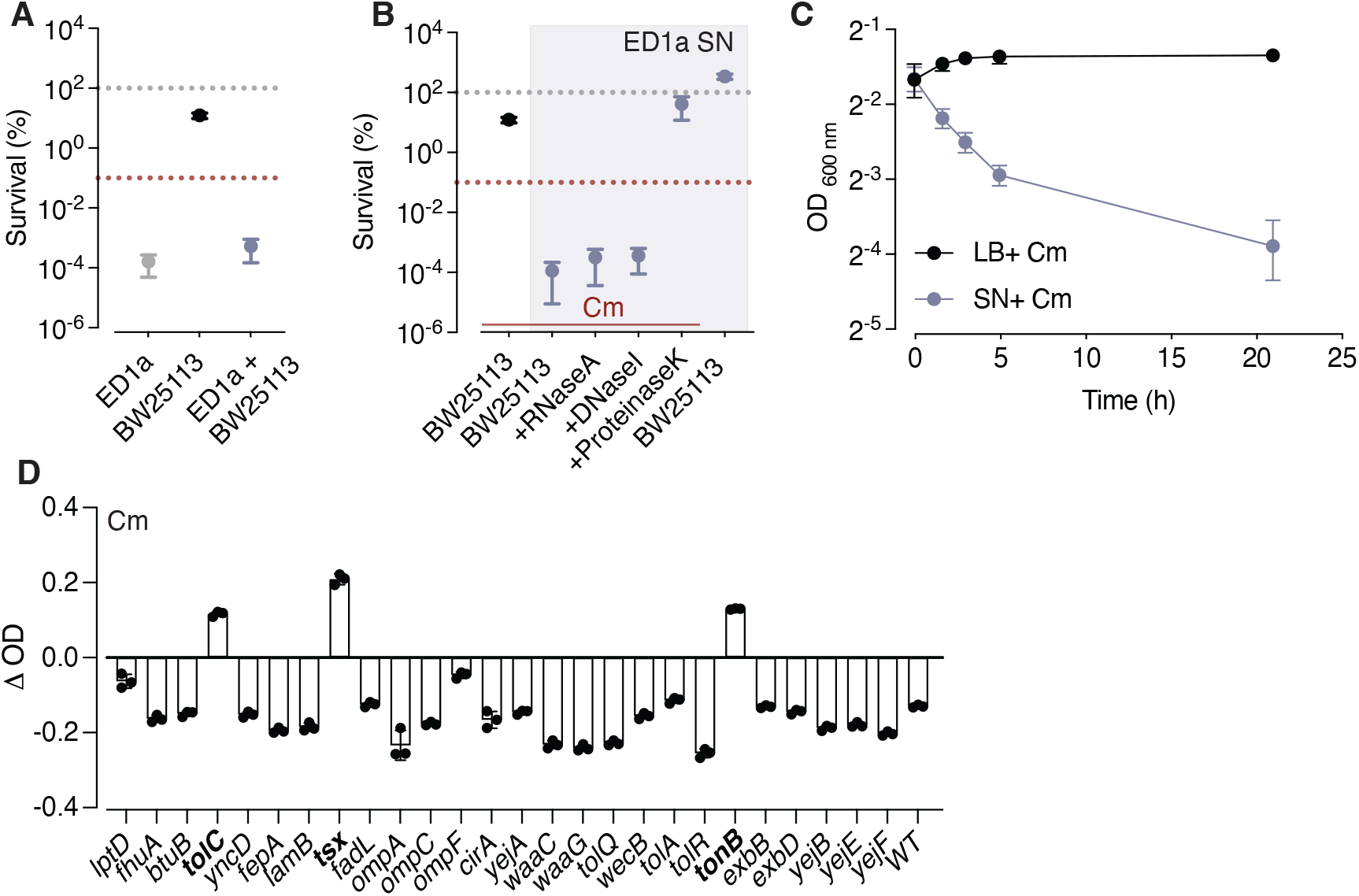
A toxic protein present in the ED1a supernatant is responsible for the killing of BW25113. All survival graphs represent the percentage survival (i.e. the number of CFU after 5 h of treatment divided by the number of CFU at time 0, multiplied by 100) and show the mean±SD of 3 independent experiments. BW25113 samples are black, ED1a, grey, and BW25113 treated with ED1a supernatant (SN) are blue. The grey dotted line corresponds to 100% survival, while the red one is 0.1%, considered the bactericidal threshold. **A.** ED1a kills BW25113 upon chloramphenicol treatment. Survival of ED1a and BW25113 alone or in combination after a 5 hour-chloramphenicol treatment. **B.** The killing activity is present in ED1a supernatant and eliminated by proteinase K. ED1a supernatant only (no chloramphenicol) does not have an impact on BW25113 survival. The highlighted zone represents samples treated with ED1a SN whereas the red line represents samples treated with chloramphenicol. Cm, chloramphenicol; SN, supernatant. **C.** BW25113 lyses upon exposure to ED1a SN and chloramphenicol (blue), but not with chloramphenicol only (black). This graph represents the mean±SD of 3 independent experiments. **D.** Deletions of *tolC, tsx* and *tonB* protect BW25113 from lysis upon exposure to ED1a supernatant and chloramphenicol. This graph shows the ΔOD (i.e. OD after 20 h of treatment – OD at the beginning) as the mean±SD of 3 independent experiments. WT is the BW25113 WT control.

### ED1a encodes bacteriocins on uncharacterized small plasmids

To identify the toxic protein, we produced and concentrated ED1a supernatant and fractionated it through a size-exclusion chromatography column. After testing individual elution fractions for activity (**Fig. 2A**), we performed proteomics analyses by mass spectrometry on five fractions (one without, two with strong and two with intermediate activity) (**Fig. 2A**). The proteome of ED1a comprises ∼5400 proteins while the one of BW25113, ∼4400, with 4000 common proteins between the two strains. In addition, ED1a carries a well-described plasmid, pECED1a, harboring 126 genes (**Fig. 2B**). From the proteins identified by mass spectrometry (**Table S3**), we eliminated (i) the proteins that were both detected in the active and inactive fractions at similar levels, and (ii) the proteins that are conserved in BW25113, as we expect the toxic protein to be absent in that strain. After applying those filters, none of the identified protein remained. We therefore expanded our search, using mass spectrometry raw data, outside of the known ED1a proteome database. This led to the identification of additional proteins, including colicin K and pesticin (**Table S3**). Both are bacteriocins that are not encoded on the ED1a chromosome or pECED1a plasmid, and were therefore not part of the initial ED1a proteome database. Bacteriocins act as microbial weapons in bacterial competition. They are proteins produced by bacteria to kill or inhibit the growth of closely related bacterial strains. Colicin K is an *E. coli*-specific bacteriocin that kills competitors by forming pores in their inner membrane, which leads to loss of membrane potential and eventually cell death^24^, while pesticin is a bacteriocin first identified in *Yersinia Pestis* which induces cell lysis by degrading peptidoglycan (muramidase activity)^25^.

**Figure 2.**
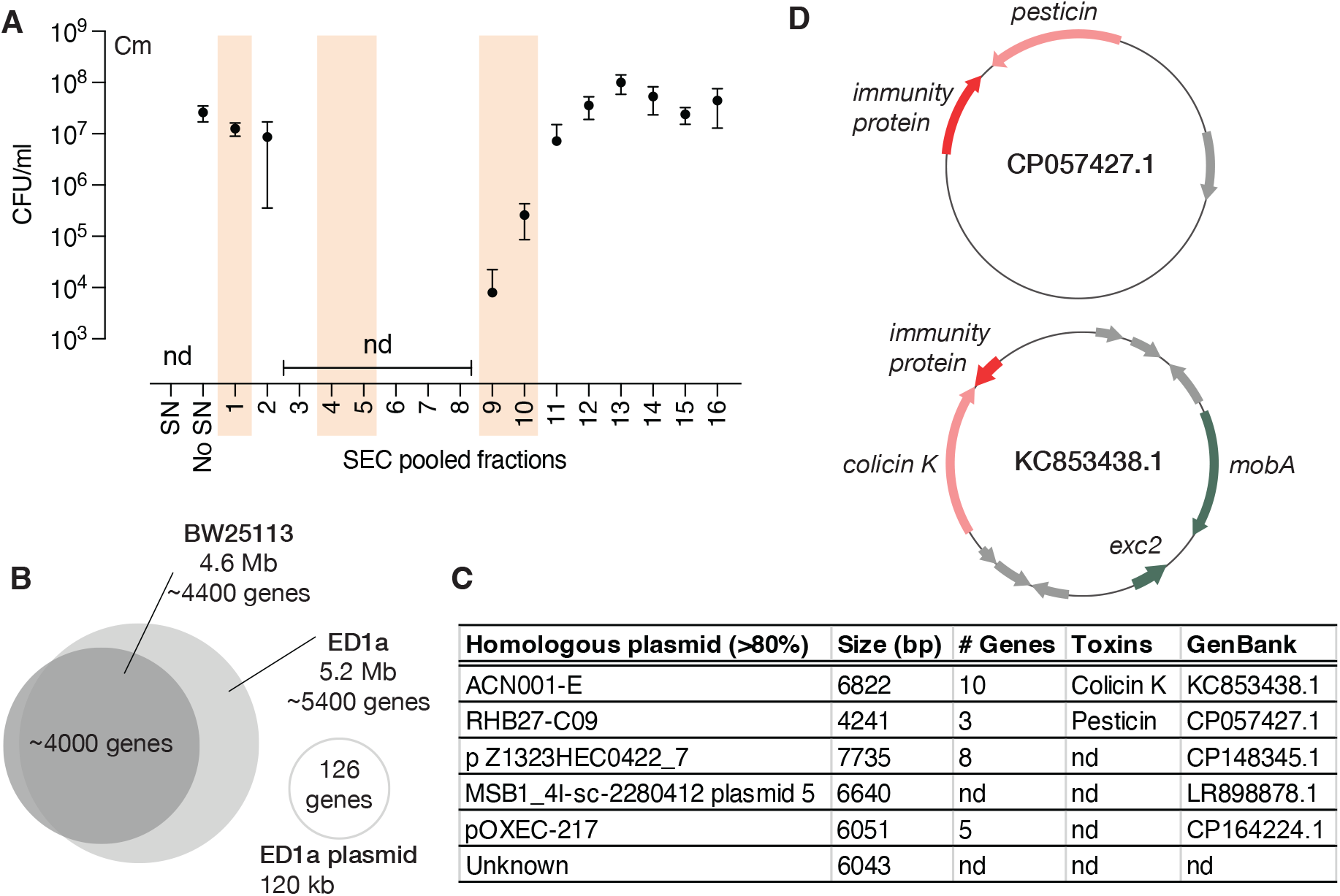
ED1a supernatant contains colicin K expressed from an unreported small plasmid. **A.** Size-exclusion chromatography elution fractions show killing of BW25113. BW25113 was exposed to SEC elution fractions combined with chloramphenicol. This graph shows the mean±SD of 3 independent experiments and the survival is assessed in colony forming units (CFU) per ml. Cm, chloramphenicol; SEC, size-exclusion chromatography; nd, not detected; SN, supernatant. The fractions highlighted in pink were further analysed by mass spectrometry. **B.** Schematic of the genetic contents of BW25113 and ED1a. **C.** ED1a carries small plasmids. The small plasmids identified in ED1a are homologous (>80%) to the ones presented in this table; nd, not defined. **D.** The identified plasmids carrying colicin K and pesticin are represented. The colicin K and pesticin genes are represented in pink, their immunity proteins in red, other annotated genes in green and predicted genes of unknown function in grey.

As bacteriocins are typically expressed from small plasmids, we next explored whether ED1a was harboring any additional plasmid, that had not been described or detected before, by performing plasmid isolation (See Methods) and sequencing. Unexpectedly, we identified six different small plasmids (**Fig. 2C**), two of which indeed encoded either the colicin K or the pesticin, along with their respective immunity proteins (**Fig. 2C-D**).

### Colicin K is responsible for the killing of BW25113 during chloramphenicol treatment

To test whether colicin K or pesticin was responsible for the killing of BW25113 upon chloramphenicol treatment, we overexpressed their respective immunity proteins in BW25113 and exposed these strains to ED1a supernatant and chloramphenicol (**Fig. 3A**). While the pesticin immunity protein did not protect BW25113 from killing, overexpression of the colicin K immunity protein (colKIP) did (**Fig. 3A**). As these immunity proteins are known to be specific, this strongly suggests that colicin K is the toxic protein present in the ED1a supernatant. We next sought to cure the small plasmids from ED1a, to get rid of colicin K production. To do so, we expressed the DdmDE plasmid defense system^18^ in ED1a. The DdmDE system is originally a defense mechanism against foreign DNA, including plasmids, that degrades small plasmids non-specifically^18^. Cloning and expressing this system in ED1a led to the curation of all small plasmids (**Fig. 3B**), but not the large one (data not shown). The supernatant of the ED1a strain lacking the plasmids was not toxic to BW25113, while re-introducing colicin K on a plasmid in ED1a restored supernatant toxicity (**Fig. 3C**). As a final step, we sought to confirm that we could detect colicin K within BW25113 cells upon supernatant exposure. We exposed BW25113 to ED1a supernatant for 30 minutes (without antibiotics) and performed proteomics on the BW25113 cells. We specifically looked for proteins that were (i) only present in BW25113 after treatment, but not before and (ii) only encoded in ED1a, but not in BW25113. This led to the identification of 2 proteins, Peptidase S6 (annotated as an IgA endopeptidase from phage origin, that is not expected to have activity on bacterial cells) and colicin K (**Table S4**), while pesticin was not detected inside BW25113 after treatment, as expected. Taken together, these results show that colicin K is the toxic protein killing BW25113 upon protein synthesis inhibitors treatment.

**Figure 3.**
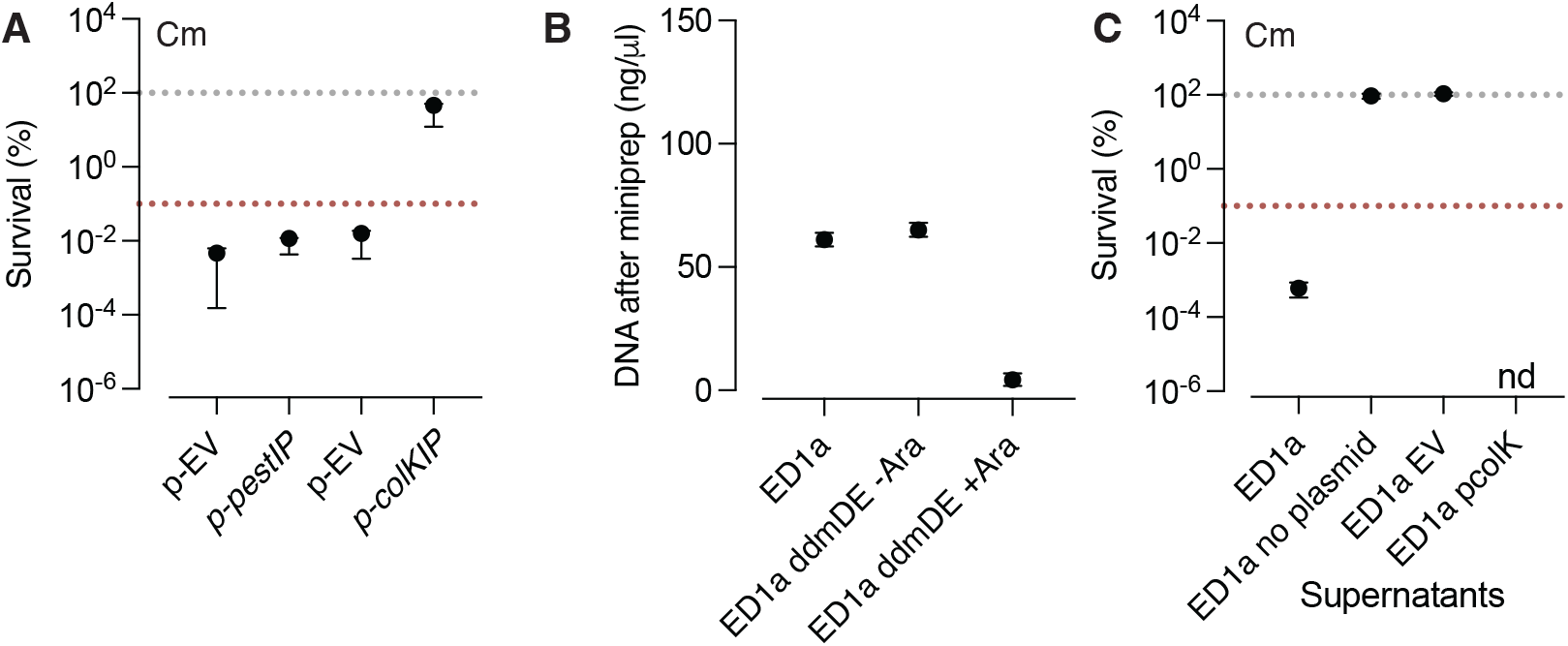
Colicin K kills BW25113 upon protein synthesis inhibitors treatment. All survival graphs represent the percentage survival (i.e. the number of CFU after 5 h of treatment divided by the number of CFU at time 0, multiplied by 100) and show the mean±SD of 3 independent experiments. The grey dotted line corresponds to 100% survival, while the red one is 0.1%, considered the bactericidal threshold. **A.** Colicin K immunity protein protects BW25113 from killing. This graph represents the survival of BW25113 that either carries an empty vector (p-EV) or a vector for the overexpression of the pesticin immunity protein (p-pestIP) or the colicin K immunity protein (p-colKIP). While the overexpression of pestIP does not improve BW25113 survival, the overexpression of colKIP does. **B.** Expression of DdmDE removes small plasmids from ED1a. This graph represents the amount of DNA left after a miniprep of WT ED1a, and ED1a carrying the DdmDE system, expressed or not with arabinose. Upon arabinose induction, small plasmids are effectively removed from ED1a. **C.** Colicin K is responsible for the toxicity of the ED1a supernatant. This graph represents the survival of BW25113 either exposed to the supernatant of WT ED1a, ED1a without its small plasmids, ED1a without its small plasmids transformed with an empty vector (EV) or ED1a without its small plasmids transformed with a plasmid expressing colicin K (pcolK). nd, not detected.

### Colicin K only kills BW25113 upon antibiotic treatment

Colicin K only kills BW25113 under chloramphenicol treatment, which is surprising as BW25113 does not encode an immunity protein against this bacteriocin. We wondered whether this strain encoded another protein that could protect it from colicin K, but which could not be expressed during protein synthesis inhibition. To test this hypothesis, we treated BW25113 with ED1a supernatant for 30 minutes without antibiotic and compared the proteome before and after treatment (**Table S5**). If BW25113 encodes a protein that protects it from the colicin K, it should be upregulated in the treated samples. We identified 17 upregulated proteins (**Fig. 4A**) and tested each of their corresponding deletions for impaired survival in the presence of ED1a supernatant as compared to BW25113 WT. None of the mutants displayed any significant change in survival (**Fig. 4B**). As most of the identified proteins are typically upregulated upon entry into stationary phase, we rationalized that their upregulation was due to the exposure of BW25113 to the ED1a supernatant, which is produced at high optical density and mimics conditions of stationary phase, rather than because of exposure to colicin K.

**Figure 4.**
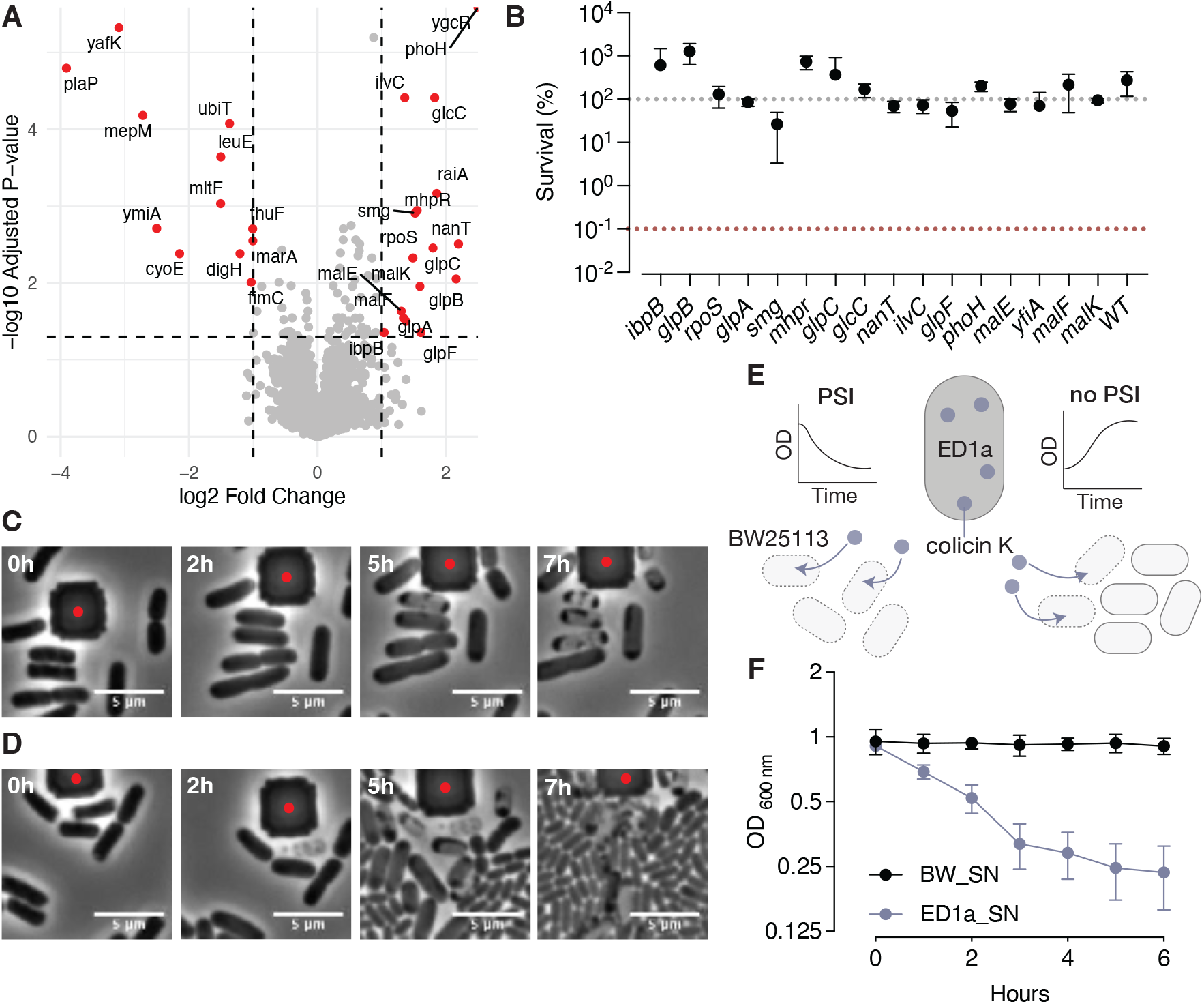
Colicin K eliminates non-growing cultures of BW25113. **A.** Exposure of BW25113 to ED1a supernatant leads to overexpression of 17 proteins, as depicted in this volano plot. Proteins in red are the ones with a log2 fold change (treated vs. non-treated) > 1 and a -log10 adjusted p-value <0.05. Results are compiled from 5 independent replicates of each condition. **B.** Deletions of overexpressed proteins do not impact BW25113 survival upon ED1a supernatant treatment. This graph shows the survival of each deletion mutant upon exposure to ED1a supernatant without antibiotic. Deletions do no show a significant negative impact on BW25113 cells. **C-D.** BW25113 cells die upon colicin K exposure with and without chloramphenicol exposure. Phase contrast time-lapse microscopy of BW25113 exposed to ED1a supernatant with (**C**) or without (**D**) chloramphenicol. Each panel shows different timepoints of the same area, with a scale bar of 5 µm. The corresponding movies are available in the supplementary material (**Supplementary Movies 1-2**). **E.** Model of the killing of BW25113 by colicin K combined or not with protein synthesis inhibitors (PSI). **F.** Stationary BW25113 lyse when exposed to a concentrated supernatant of ED1a, without chloramphenicol. This graph shows the mean±SD of 3 independent experiments of the decrease in OD observed after addition of concentrated ED1a supernatant (ED1a_SN, blue) or BW25113 supernatant (BW_SN, black) to stationary phase BW25113 cells.

Another hypothesis to explain that BW25113 survives colicin K under normal condition is that the bacteriocin is not concentrated enough to have a visible impact on growing bulk cultures (i.e. it kills a few cells, but the effect is masked by the non-affected cells that are still growing). To test this hypothesis, we performed time-lapse microscopy, using microfluidics, to observe BW25113 cells treated with ED1a supernatant in the presence or absence of chloramphenicol. As expected, treatment with supernatant and chloramphenicol led to strong lysis and cell death in BW25113, with dead cells displaying a patchy phenotype (**Fig. 4C – Supplementary Movie 1**). This is surprising as it does not look like the typical effect expected from pore-forming toxins, suggesting that colicin K might have a slightly different mode-of-action on this strain. Interestingly, when BW25113 was treated with ED1a supernatant without any antibiotic, we could still detect cells dying with a similar phenotype, but the rest of the cells were growing (**Fig. 4D – Supplementary Movie 2**). This observation suggests that colicin K kills BW25113 but is not concentrated enough to kill the whole population, especially as it is growing. This means that in a growing culture, the effect of colicin K is barely visible, while it has a way stronger impact on a non-growing population (**Fig. 4E**). To confirm our hypothesis, we tested the effect of concentrated ED1a supernatant on a non-growing stationary phase culture of BW25113. As expected, we observed that non-growing BW25113 cells were lysing, confirming that colicin K killing activity affects and becomes visible on non-growing BW25113 cells, even in the absence of a protein synthesis inhibitor (**Fig. 4F**).

### Antibiotic treatment influences bacterial competition

Many bacterial strains carry bacteriocins, suggesting that the effect we observe might be conserved across diverse strains or specied. To investigate how broadly competition is modulated by antibiotic treatment, we decided to work with a library of 1,085 natural isolates of *E. colis*. To this end, we collected their supernatants and assessed their effects on BW25113 in the presence of chloramphenicol (**Fig. 5A**). We specifically looked for supernatants that caused a decrease in optical density (>30% decrease) under chloramphenicol treatment, indicative of toxicity and cell lysis (**Fig. 5A**). Of the 1,085 supernatants tested, 108 exhibited toxic activity against BW25113 under these conditions (**Fig. 5A**). As high-throughput screens sometimes lead to false positives, we repeated the test with lower throughput for the 108 toxic supernatants and confirmed strong lysis activity for 74 of them (**Fig. 5B – red dots**), none of which showed lysis in the absence of chloramphenicol (data not shown). To explore whether lysis was driven by a similar mechanism as the one with colicin K, we tested whether the chloramphenicol-dependent killing activity of these supernatants depended on the presence of a toxic protein, as it was the case for ED1a. We exposed the 74 toxic supernatants to proteinase K, which suppressed the lytic activity for 71 out of the 74 samples (**Fig. 5B – grey dots**), suggesting that toxin-mediated bacterial killing is a common mechanism that is strongly influenced by antibiotic presence. Overall, our results show that the antibiotic-dependent protein-mediated toxicity is quite prevalent between competing bacteria.

**Figure 5.**
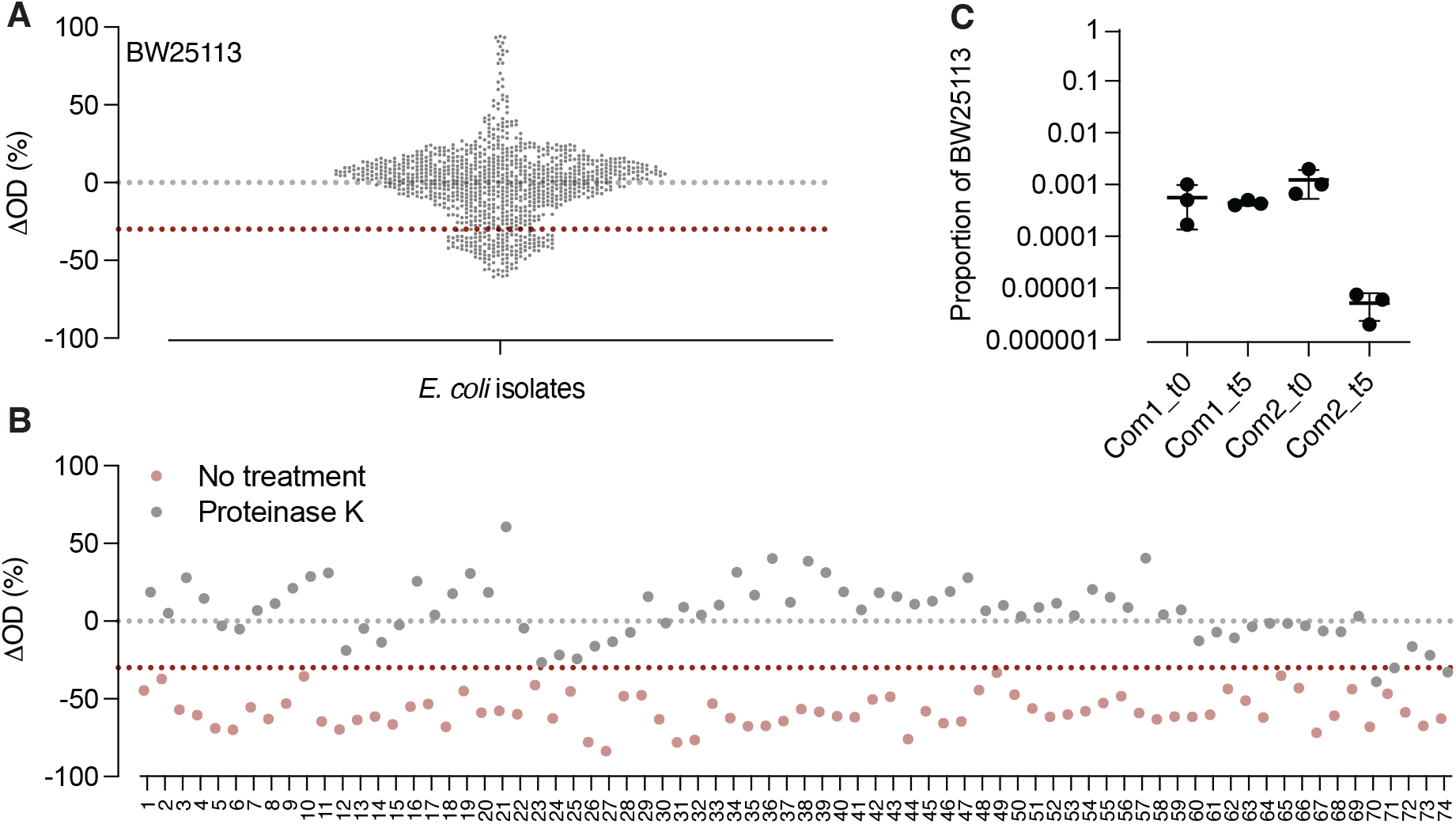
Antibiotic-dependent toxin killing of BW25113 is conserved among *E. coli* natural isolates and impacts bacterial communities. **A.** BW25113 is killed by 108 supernatants out of the 1085 tested *E. coli* isolates (threshold >30% lysis). This graph shows the mean of 2 independent experiments. Percentage lysis is calculated as (OD_end_ − OD_beginning_) / OD_beginning_ × 100 of BW25113 cultures before and after addition of supernatant combined with chloramphenicol (Cm) for 24h. **B.** 74 supernatants were confirmed to kill BW25113 upon chloramphenicol treatment, 71 of which harbor a toxic protein. This graph shows the mean of 2 independent experiments. Percentage lysis is calculated as (OD_end_ − OD_beginning_) / OD_beginning_ × 100 of BW25113 cultures before and after addition of supernatant (pre-treated or not with proteinase K) combined with chloramphenicol (Cm) for 24h. Correspondance to strain names is included in Supplementary Table 6. **C.** BW25113 is eliminated from a community that also harbors ED1a upon chloramphenicol treatment. This graph shows mean±SD of 3 independent experiments, and depict the proportion of BW25113 cells as compared to the whole community before (t0) and after (t5) chloramphenicol treatment. Com_1 is *community 1* and harbors 11 diverse gut microbes and BW25113 as the sole *E. coli* strain. Com_2 is *community 2* and harbors 11 diverse gut microbes and BW25113+ED1a as *E. coli* strains.

In parallel, we explored how this antibiotic-dependent toxicity impacted bacterial communities. We assembled a synthetic community of 11 diverse gut microbial members that also included either BW25113 only (community 1) or both BW25113 and ED1a strains (community 2). The communities were exposed to chloramphenicol for 5h, and CFU were counted before and after treatment. By counting the *E. colis* remaining in each community, we observed that in the community harboring BW25113 as the sole *E. coli* strain (community 1), its counts remained stable before and after treatment, as expected for a treatment with a *bacteriostatic* antibiotic. However, in the community harboring both ED1a and BW25113 (community 2), BW25113 counts were strongly decreased (∼ 3log_10_) during treatment (**Fig. 5C**), confirming that the effect of ED1a and colicin K is conserved in a community setting. This discovery is important as it highlights an unexpected effect of antibiotic treatment and provides another layer of explanation on how protein synthesis inhibitors induce strong dysbiosis in gut bacterial communities while not always showing strong impact on individual strains^9–13^.

## Discussion

Antibiotics are traditionally defined by their intrinsic activity on essential bacterial processes, yet our work reveals that the outcome of antibiotic treatment can be strongly shaped by the genetic context of the target strain and its neighbours. Here, we show that bacteriostatic protein synthesis inhibitors trigger the elimination of BW25113 through colicin K, a bacteriocin encoded on a small plasmid carried by ED1a and normally deployed in inter-strain competition. These findings redefine antibiotics not as universally bacteriostatic or bactericidal agents, but as modulators whose ultimate effects can be conditioned by accessory genes such as bacteriocins, toxin–antitoxin systems, or phage defence modules^26^. This perspective helps explain why antibiotics can exert disproportionate, strain-specific killing in microbial communities that would not be predicted from MIC-based classifications alone.

We found that chloramphenicol synergizes with colicin K secreted by ED1a to eradicate BW25113, whereas colicin K alone has little effect on a BW25113 culture, and chloramphenicol alone exerts only its classical bacteriostatic action. This synergy is not an idiosyncrasy of the ED1a–BW25113 pair: screening a library of 1,085 natural *E. coli* isolates, we found that antibiotic-induced, toxin-mediated killing is a widely conserved interaction, suggesting that this mechanism operates broadly across *E. coli* populations and likely extends to other bacteriocin-producing species. Moreover, we show that this interaction is not confined to simple pairwise co-cultures: in a synthetic community of 12 gut bacterial members that included both ED1a and BW25113, exposure to a protein synthesis inhibitor was sufficient to drive clearance of BW25113, demonstrating that colicin-antibiotic synergy can shape strain-level outcomes even in the presence of a diverse competing microbiota. Together, these findings show that antibiotics can reconfigure competitive landscapes at both the pairwise and community level, with important consequences for microbiome stability in environments such as the gut or aquatic ecosystems that are regularly exposed to antibiotics.

Finally, our results highlight the underestimated role of small plasmids in natural bacterial isolates. We were surprised to detect six small plasmids in ED1a that had not been previously reported, even in studies examining the probiotic potential of ED1a^27^, likely a consequence of relying on short-read genome sequencing. This underscores how limited our knowledge of natural isolates remains, even when their genome sequences are ostensibly available. Our work shows that despite being frequently overlooked and poorly annotated, these mobile elements encode genes with profound effects on bacterial physiology and antibiotic responses, effects that, as our conservation analysis suggests, are not restricted to a single strain pair but recur across natural populations.

In summary, we show that the fate of a bacterium under antibiotic stress is determined not solely by drug-target interactions, but also by the genetic arsenal carried by the bacterium itself and by its neighbours, and that this phenomenon is broadly conserved rather than a peculiarity of one strain pair. This insight opens a path toward predicting antibiotic outcomes from genome content and community composition, and suggests that optimizing antimicrobial therapies will require moving beyond target-based classifications toward community- and genome-aware frameworks.

## Legend (supplementary material)

**Supplementary Movie 1.** BW25113 cells die upon colicin K exposure when combined with chloramphenicol. Phase contrast time-lapse microscopy of BW25113 exposed to ED1a supernatant with chloramphenicol.

**Supplementary Movie 2.** BW25113 cells also die upon colicin K exposure when not combined with chloramphenicol, but most cells are not affected and grow. Phase contrast time-lapse microscopy of BW25113 exposed to ED1a supernatant without chloramphenicol.

**Table S1** – list of strains used in this study

**Table S2** – list of plasmids used in this study

**Table S3** – proteomics data obtained from studying the SEC fractions leading to killing of BW25113

**Table S4** - proteomics data obtained from studying the proteins present inside BW25113 after treatment with ED1a supernatant

**Table S5** - proteomics data obtained from studying the proteins overexpressed in BW25113 after exposure to ED1a supernatant

**Table S6** – correspondance to strain names of Figure 5B.

## Acknowledgements

We thank Nassos Typas for providing valuable advice, strains and support to this project. Marco Galardini, Nicolai Karcher and Alexandra Koumoutsi for their help with the E. coli library genomes. We acknowledge Melanie Blokesch for support of C.V., provision of materials, and scientific discussion regarding the DdmDE-mediated plasmid chase experiment. Finally, we thank the EPFL proteomics core facility (specifically Maria Pavlou and Mathilde Willemin) and PTPSP EPFL core facility (Florence Pojer and Yoan Duhoo) for their help and valuable support.

## Conflict of interest

There is no conflict of interest.

## Authors contribution

This study was conceived, designed and supervised by CVG. Most experiments were designed and performed by VR, JG, KM, KV, YA, CS and CVG. CV supported us with the microfluidics experiments. Data interpretation was performed by VR, JG, KM and CVG. CVG wrote the manuscript and designed the figures with feedback from all authors. All authors approved the final version for publication. This work was supported by the SNSF Starting Grant (TMSGI3_226334/1/SNSF Starting Grant) for VR and JG, and by the SNSF Project Funding (10002165/SNF) for KM.

## Notes

### Competing Interest Statement

The authors have declared no competing interest.

